# The HIV-1 Vpu transmembrane domain topology and formation of a hydrophobic interface with BST-2 are critical for Vpu-mediated BST-2 downregulation

**DOI:** 10.1101/2020.06.28.176289

**Authors:** Nabab Khan, Siladitya Padhi, Paresh Patel, U. Deva Priyakumar, Shahid Jameel

## Abstract

Viruses belonging to the M group of human immunodeficiency virus (HIV-1) are the most virulent among the four HIV-1 groups. One factor that distinguishes the M group HIV-1 from others is Vpu, a membrane localized accessory protein, which promotes the release of virions by neutralizing the antiviral host cell protein BST-2. To investigate if this activity is determined by the topology of Vpu or by conserved amino acid residues, we prepared chimeric forms of Vpu by replacing its transmembrane domain with those from its topological homologs. Although the chimeric Vpu proteins downregulated BST-2, these substantially reduced virus production as well. Molecular modeling studies on Vpu from different HIV-1 groups and the chimeric Vpu proteins showed that shape and the availability of a hydrophobic interface are more important for BST-2 antagonism than conservation of the amino acid sequence. Our data suggest that the HIV-1 Vpu-M protein has evolved topologically to interact with BST-2, and that the Vpu/BST-2 interface can be exploited as a target to limit HIV-1 replication.

## Introduction

The human immunodeficiency virus type 1 (HIV-1) has been classified into four different groups, namely M, N, O and P, of which the M group is responsible for a majority of global infections [1,2]. The restriction factor BST-2 (also called Tetherin or CD317) binds HIV-1 and prevents its release from the cell surface, resulting in lower levels of extracellular virions [3, 4]. The human and simian immunodeficiency viruses have evolved different mechanisms for counteracting BST-2. While SIVcpz and SIVgor use the viral Nef accessory protein for this purpose [5–7], HIV-1 encodes the viral protein U (Vpu) to overcome BST-2 [3]. Among the HIV-1 strains, viruses that belong to the M and N groups have evolved Vpu proteins with anti-BST-2 activity, but Vpu from HIV-1 O and P groups do not possess this activity [8, 9].

Vpu is a 16-kDa accessory protein of HIV-1 [10–12], which promotes degradation of the CD4 receptor, release of new virions and evasion from host immune responses [13–16]. Upon being expressed intracellularly, Vpu localizes to the plasma membrane, endoplasmic reticulum (ER) membrane and transgolgi network (TGN) [17–19]. It has two distinct structural domains with different biological activities. The N-terminal transmembrane (TM) α-helical domain is required for the budding of new virions from infected cells, and for cation-selective ion channel activity in the plasma membrane. The cytoplasmic domain is involved in CD4 degradation [20–22], and consists of two a-helices that are linked by a loop, which contains Serine-52 and −56 that can be phosphorylated [23, 24]. These phosphoserines are involved in the ubiquitination and proteosomal degradation of BST-2 and CD4 through the assembly of a b-TrCP/Skp1/Cul1 complex [25–27]. Moreover, Vpu promotes virus release by redirecting BST-2 from the cell surface towards lysosomal degradation, and by diverting CD4 from the ER membrane towards the ER-associated degradation (ERAD) pathway [27]. Vpu also enhances Gag transport to the virion assembly site by weakening its interaction with the small glutamine-rich tetratricopeptide repeat-containing protein alpha (SGTA) [28, 29].

BST-2 is a single-pass type-II transmembrane protein that is localized to cholesterol-rich domains of the plasma membrane and TGN [30]. It is composed of an N-terminal transmembrane domain and an extracellular domain that is heavily glycosylated and has a glycophosphatidylinositol (GPI) anchor [31]. During virus release, BST-2 captures virions on the cell surface via its transmembrane domain (TMD) and GPI anchor [32]. But since Vpu removes BST-2 from the cell membrane, it helps overcome this restriction [33]. The helical TMDs of Vpu and BST-2 interact with each other in an anti-parallel orientation [34]. The Ala14, Ala18 and Trp22 residues within the Vpu TMD are reported to be involved in BST-2 downregulation from the plasma membrane via interaction with specific residues within the BST-2 TMD [20]. However, the interaction between BST-2 and Vpu is lost when the polarity of the Vpu TMD is changed, even in the presence of these critical residues [35], suggesting that Ala14, Ala18 and Trp22 by themselves are not the major determinants for this function. The Vpu TMD is highly hydrophobic with alanines that are spaced four residues apart (Ala10, 14, 18), and these show a high propensity for helix stabilization.

Since changing the polarity of the Vpu TMD inhibits helix-helix interaction, topology could be important for the interaction. To investigate this, we computationally identified two topological homologues of the Vpu TMD, and then synthesized recombinant forms of Vpu by replacing its TMD with those of the homologues. The recombinant forms reduced BST-2 levels by degrading it through the proteasomal pathway, thus confirming the functional role of topology in this interaction. These results are further supported by molecular modeling studies, which show that topology of the Vpu TMD and the availability of a hydrophobic interface on Vpu and BST-2 are crucial determinants of this interaction. However, unlike Vpu, the recombinant TMD-substituted Vpu proteins inhibit HIV-1 replication. This opens up the possibility of targeting this interface as an antiviral target.

## Results

### Expression and characterization of recombinant Vpu

To identify potential homologs of the Vpu TMD, sequences of 510 α-bitopic proteins and 6751 α-polytopic proteins were obtained from the Topology Data Bank of Transmembrane Proteins (TOPDB) [36]. Additionally, 850 transmembrane segments were obtained from the Membrane Protein Topology Database (MPtopo) [37]. **Table 1** shows the sequences that had a score greater than 25 and more than 70% structural homology with the Vpu TMD (see Methods section). The choice of TMDs for further analyses was essentially random, with the premise that if topological effects were important, most of these homologs should work. The only choices we made were to choose one bacterial and one human sequence, and one from a bitopic protein and another from a polytopic protein. Consequently, TMDs from proteins with accession IDs AB00681 and AP00387 were chosen to investigate their topological homology with Vpu and functional effects on BST-2. AB00681 is the human Asialoglycoprotein receptor 1 (also called hepatic lectin H1), which is a single-pass transmembrane protein that is expressed in hepatic parenchymal cells [38]. This receptor recognizes terminal galactose and N-acetylgalactosamine units on glycoprotein ligands [39]. AP00387 is the E. coli Acriflavine resistance protein B, a multi-pass transmembrane protein that is involved in drug efflux [40–42].

**Table 1.**
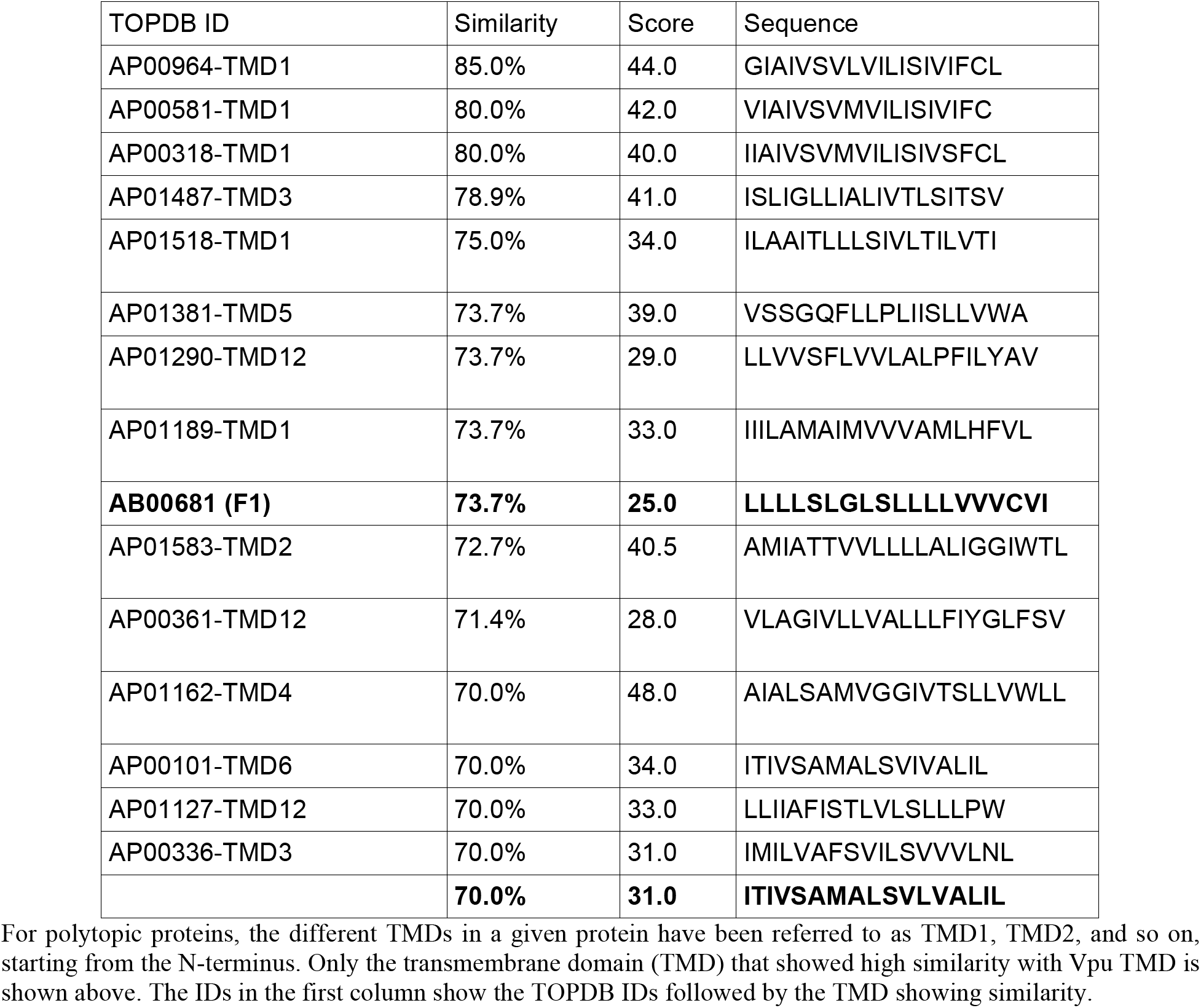
Sequences from TOPBD showing >70% similarity with the Vpu-M transmembrane domain (TMD)

We replaced the TMD of Vpu-M with those from AB00681 and AP00387; these are called F1 and F2, respectively (**Fig. 1A**). The Vpu-M, F1 and F2 proteins were expressed as fusions to the enhanced green fluorescent protein (EGFP). While both proteins showed expression in transfected cells, as evidenced by Western blotting (Fig. 1B) and EGFP fluorescence (Fig. 1C), F1-EGFP was expressed at high levels but F2-EGFP showed poor expression. Vpu trafficking to intracellular and plasma membranes is important for its BST-2 antagonism and virus release from infected cells [17–19]. We therefore checked the membrane localization of F1 and F2 by staining transfected TZM-bl cells (a HeLa cell derived HIV-permissive cell line) with the membrane tracker dye PKH26GL. The distribution of F1 and F2 proteins and their colocalization with PKH26GL was similar to that of Vpu-M (**Fig. S1**).

**Figure 1.**
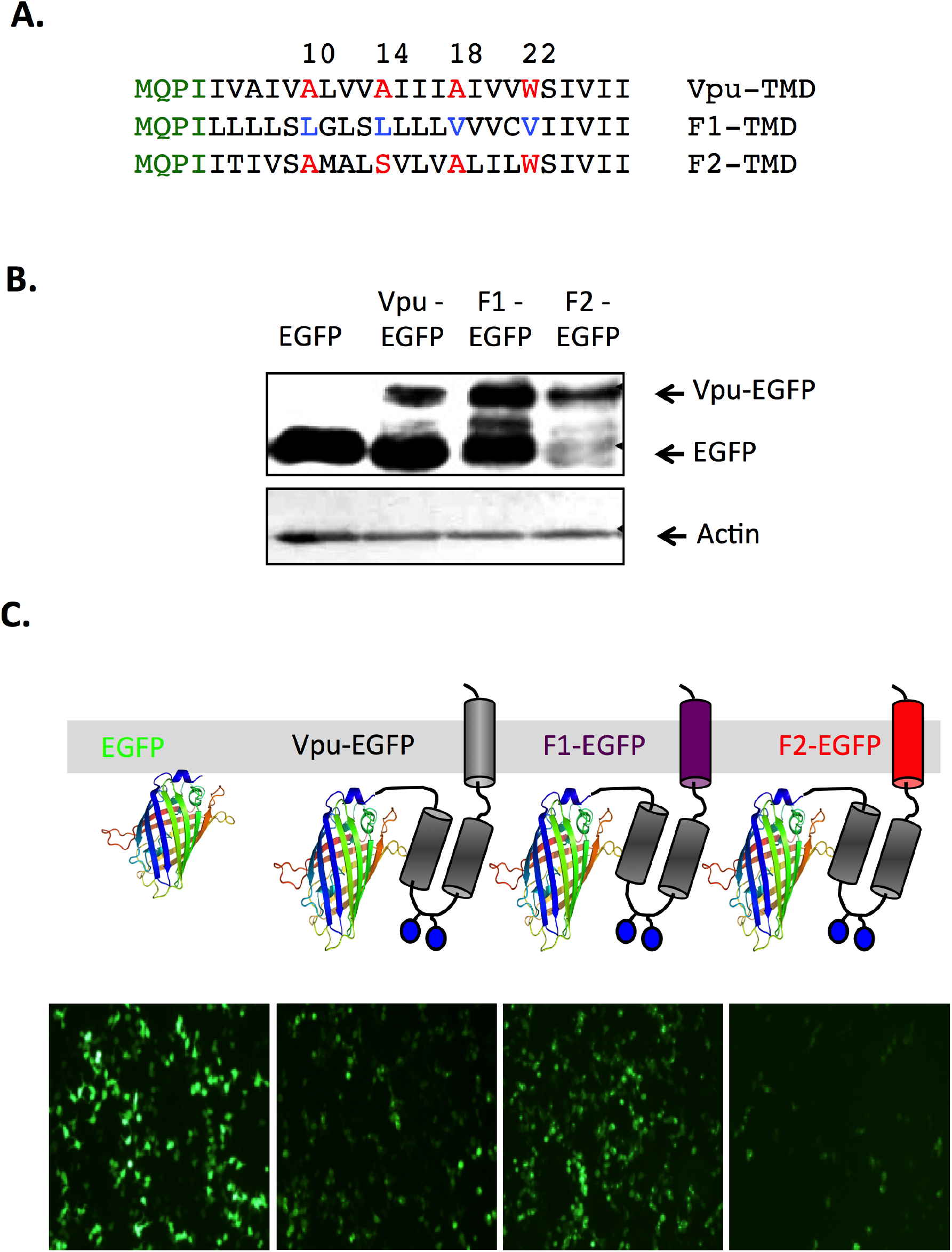
Recombinant forms of Vpu. (A) Transmembrane domain (TMD) sequences of Vpu-M, F1 and F2. The N-terminal region is shown in green, the A10(X3)A14(X3)A18(X3)W22 motif is shown in red and the hydrophobic residues replacing this motif in F1 are shown in blue. (B) Expression of EGFP-tagged Vpu-M, F1 and F2 in HEK293T cells. Western blot was developed with anti-GFP antibodies and Actin was used as a loading control. (C) Schematic of the EGFP-tagged Vpu-M, F1 and F2 proteins and expression in HEK293T cells visualized for EGFP fluorescence by confocal microscopy.

### Vpu and its homologs interact with BST-2

The Vpu-M, F1 or F2 proteins were expressed in TZM-bl cells, which endogenously express BST-2. All three proteins were similarly distributed in the cells and showed colocalization with BST-2 (**Fig. 2A**). We then cotransfected HEK293T cells, which are negative for BST-2, to express EGFP-tagged Vpu, F1 or F2 and HA-tagged BST-2. When the cell lysates were immunoprecipitated with mouse anti-GFP and the precipitates Western blotted with rabbit anti-HA, BST-2 was found to immunoprecipitate with Vpu, F1 and F2, but not with EGFP (Fig. 2B; upper panel). Alternatively, Vpu, F1 and F2 were also found in BST-2 immunoprecipitates (**Fig. 2B**; lower panel).

**Figure 2.**
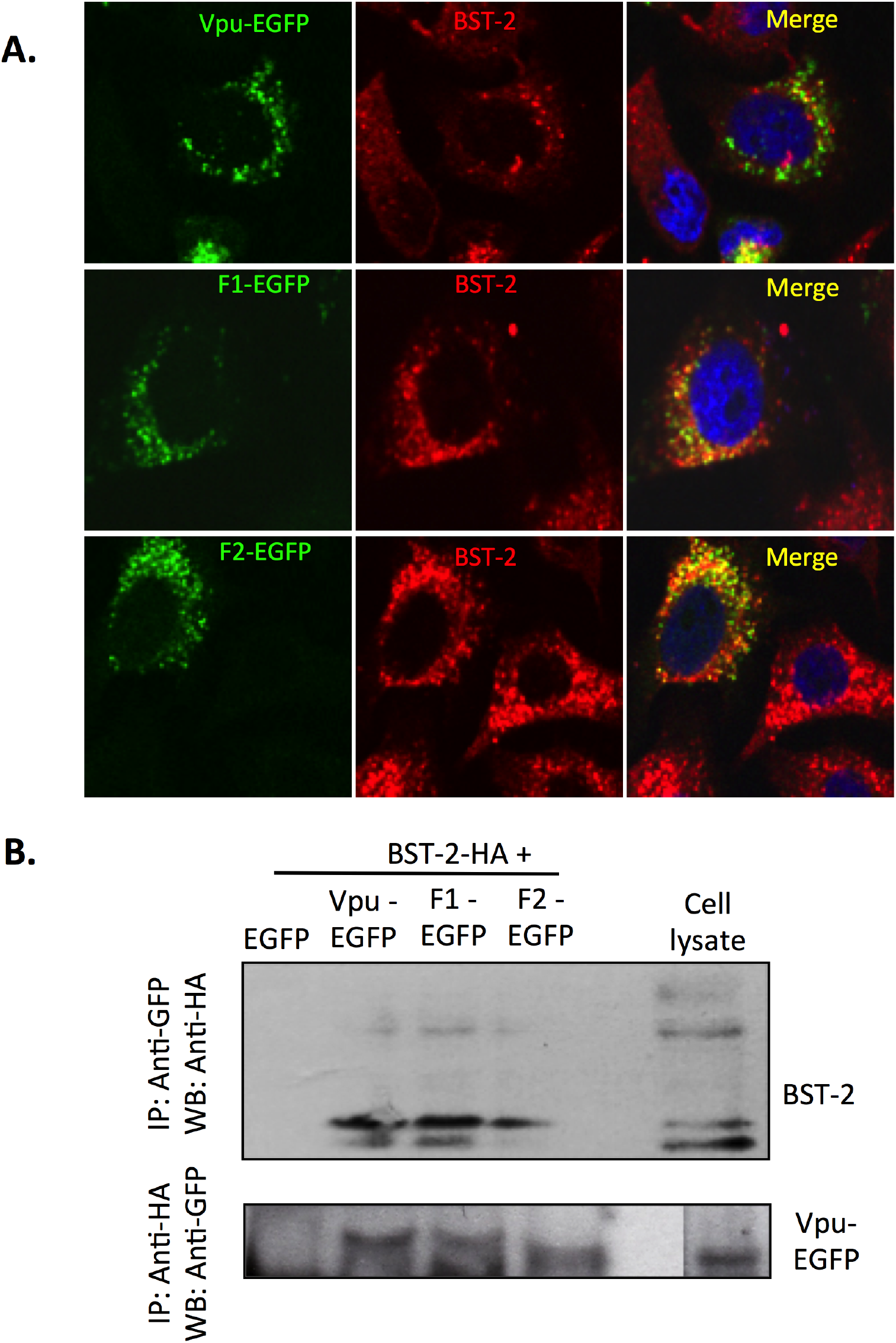
Association of Vpu with BST-2. (A) TZM-bl cells (HeLa derivative) were transfected with expression vectors for EGFP-tagged Vpu-M, F1 or F2. The cells were counter-stained with anti-BST-2 antibodies for endogenous BST-2 and visualized by confocal microscopy. (B) HEK293T cells were cotransfected with the expression vectors for HA-tagged BST-2 and EGFP-tagged Vpu-M, F1 or F2 (EGFP was used as control). The cell lysates were immunoprecipitated (IP) with anti-GFP followed by Western blotting (WB) with anti-HA (upper panel); alternatively, the lysates were IP with anti-HA followed by WB with anti-GFP (lower panel).

### The Vpu/BST-2 association is driven by hydrophobic interactions

To investigate the nature of interactions between Vpu and BST-2, monomeric forms of the TMDs of the two peptides were docked onto each other. Previous NMR studies have shown the helical TMDs of Vpu and BST-2 to pack in an anti-parallel manner [34]. Representative structures of Vpu bound to BST-2 were chosen from the docked structures such that the two helices were antiparallel and spanned the entire lipid bilayer. Thus, if the membrane extends along the horizontal plane, the N-terminus of one helix should lie in approximately the same horizontal plane as the C-terminus of the other helix, and vice-versa. This would ensure that the transmembrane regions of the two proteins lie in the same plane, allowing them to be properly positioned in the membrane. The representative structures from the docked complexes were then simulated in a fully solvated lipid bilayer environment to investigate the dynamic nature of the interaction. Figure 3 shows the contact distances between different residues of the Vpu (or its homologs) and BST-2 TMDs averaged over the simulation time. As evident from these contact maps (see also **Table S1 in Supporting Information**), the residues making close inter-helical contacts are predominantly hydrophobic. This is consistent with previous NMR studies, which show that the helix-helix association is mediated by interactions between hydrophobic residues in the Vpu and BST-2 TMDs [34].

**Figure 3.**
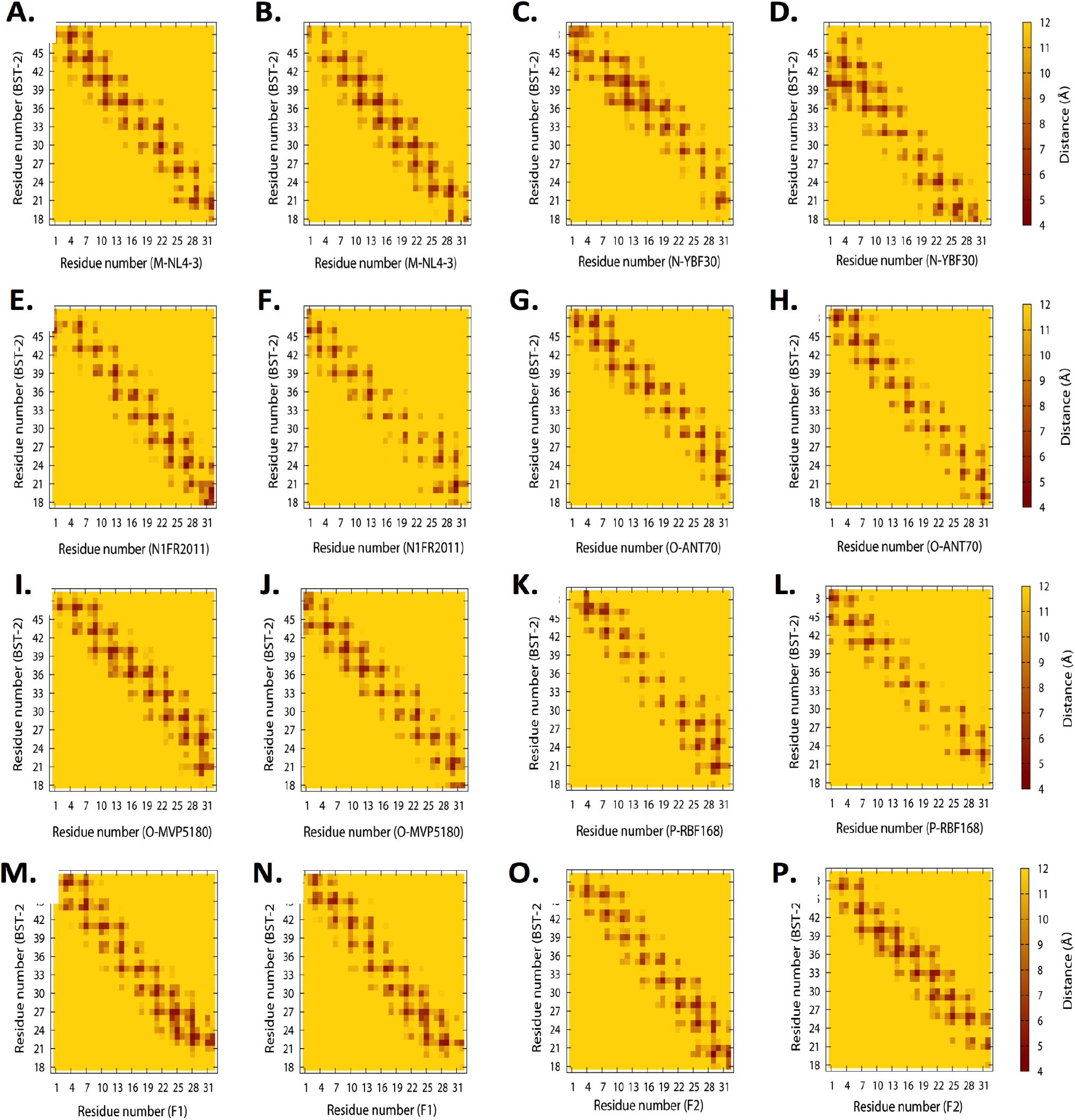
Contacts between the residues on Vpu and BST-2. Two poses were modeled for each Vpu type in complex with BST-2. The distances shown have been averaged over simulation times. (A, B) M-NL4-3; (C, D) N-YBF30; (E, F) N1FR2011; (G, H) O-ANT70; (I, J) O-MVP5180; (K, L) P-RBF168; (M, N) F1; and (O, P) F2.

The BST-2 residues that lie at the helical interface in both the model structures (referred to as “pose 1” and “pose 2”) are Ile26, Val30, Ile34, Leu37, Leu41, and Phe44. Previous experimental studies have shown Ile34, Leu37, and Leu41 to be essential for interaction with Vpu [43]. In both simulated models, all BST-2 residues that contact with Vpu are nonpolar, the only exception being Cys20 in pose 1. This suggests that the availability of a fully hydrophobic face on BST-2 is crucial for it to bind Vpu. The contact maps further reveal Vpu residues Ile4, Ile8, Leu11, Ala14, Ile15, Trp22, and Val25 to be at the interaction interface (**Fig. 3**). Of these, Ala14 and Trp22 have previously been shown to be critical for Vpu-mediated BST-2 downregulation [20].

The Vpu proteins of HIV-1 groups O and P do not downregulate BST-2 (44). As evident from the contact maps, Vpu-P does not form significant contacts with BST-2, indicating that the packing is not compact. While Vpu-O forms a number of contacts, the helix-helix interface involves more polar residues than the other proteins. The interface residues include His2, Asp5, Ser13 and Tyr30 from O-ANT70 Vpu, and Arg19, Thr45, and Lys47 from BST-2; or Asn5, Ser13, Cys16, and Tyr30 from O-MVP5180 Vpu, and Lys21 and Lys47 from BST-2. Since the Vpu types that are inactive towards BST-2 do not have a fully hydrophobic interface, it follows that the occurrence of hydrophilic residues at the interface can disrupt the interaction. This hypothesis is in good agreement with experiments on mutant forms of Vpu in which isoleucine residues at positions 15, 16, and 17 were substituted [35]. Interestingly, a mutant with the three isoleucines replaced by valines (M3IV mutant) could also downregulate BST-2 reasonably well. Since the substitutions were conservative, the protein retained a hydrophobic interface even after the mutation. However, replacement of the isoleucines with threonines (M3IT mutant) led to compromised BST-2 downregulation. Here we have identified the residue at position 15 as being a part of the interaction interface, and this mutation is expected to affect Vpu-mediated BST-2 downregulation.

A TM substitution mutant of Vpu with the Ala14→Leu, Ala18→Leu and Trp22→Ala mutations was also docked onto BST-2. However, since none of the top-scoring docking poses satisfied the two criteria described above, no simulations were performed on this complex. As far as interactions between F1/F2 and BST-2 are concerned, the TMDs of F1 and F2 packed compactly against BST-2, and the contact maps for F1 or F2 interacting with BST-2 provide insights into the interaction. A number of compact contacts with BST-2 are formed by both F1 and F2, despite the absence of Trp22 in F1, and Ala14 in both F1 and F2, suggesting that conservation of these residues is not essential for the Vpu/BST-2 interaction. While F2 has different sets of residues forming the binding interface in the two models, F1 has six residues that form contacts with BST-2 in both the models: Leu7, Leu14, Leu17, Cys21, Val25, Glu28, and Tyr29. The BST-2 residues that interact with F1 in both the models are Leu22, Leu23, Gly27, Val30, Ile34, and Thr45. The nature of the interacting residues suggests that conservation of an overall hydrophobic face is more important than conservation of a functionally important single residue. A number of close contacts are seen between BST-2 and residues in the N-terminus of Vpu. This suggests that N-terminal residues might also play a role in BST-2 downregulation, in agreement with a recent study on Vpu from HIV-1 M group subtypes B and C [45].

### Determinants of Vpu and BST-2 TMD packing

The Vpu from some of the HIV-1 groups interact with BST-2, while others do not. To investigate if this differential behavior towards BST-2 is determined by sequence or topology, both sequence- and structure-level analyses were performed on the TMD peptides. A multiple sequence alignment of the natural and recombinant Vpu types is shown in **Figure 4A**. All the BST-2-inactive Vpu types (O-ANT70, O-MVP5180, and P-RBF168) have charged residues near the N-terminus. Since the N-terminus has been suggested to play a role in BST-2 degradation and virus release [45], the inactivity of some Vpu types might be due to these charged residues. Furthermore, the possible function of the N-terminal region, hitherto unknown, might be governed by the nature of this region.

**Figure 4.**
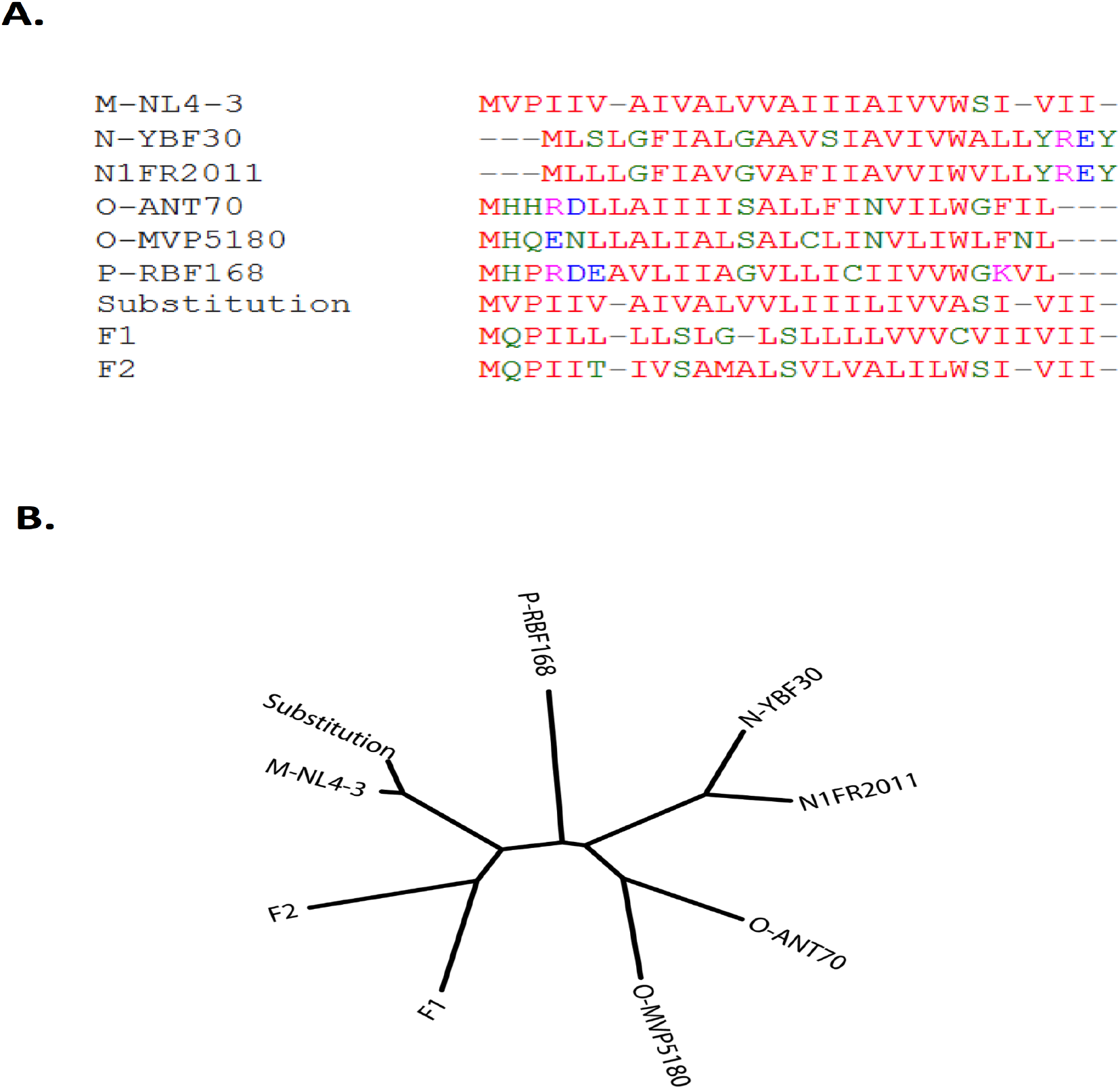
Sequence comparison of the different Vpu types. (A) Multiple sequence alignment of the TMDs of different Vpu types, including the recombinants F1 and F2. (B) A phylogenetic tree of the different Vpu types, F1 and F2 constructed on the basis of TMD sequence similarity. An unrooted neighbor-joining tree is shown.

A phylogenetic tree was constructed on the basis of sequence similarity, which showed four small clusters – M-NL4-3 and TM Substitution, F1 and F2, O-ANT70 and O-MVP5180, and N-YBF30 and N1FR2011 (**Figure 4B**). Evidently, Vpu types belonging to the same group are part of the same cluster. However, the tree suggests that there is no clear correlation between sequence conservation and BST-2 activity. Although Vpu-M and the TM substitution are in the same cluster and have good sequence homology, they differ in their ability to interact with and restrict BST-2. On the other hand, Vpu-M and Vpu-N show similar activity towards BST-2 despite differences in the primary sequence. The anti-BST-2 activity of Vpu, therefore, is determined by factors other than the sequence of the protein.

To understand how the shape of the helix varies as one moves along the helical axis from one end to the other, the side chain solvent accessible surface area (SASA) was calculated residue-by-residue and compared with that of Vpu-M (**Fig. 5**). Unlike M-NL4-3, the N-YBF30 and N1FR2011 Vpu have their surface protruding inward at several places due to the presence of glycine residues. While N-YBF30 has a bulky Phe residue near the N-terminus (**Fig. 5A**), N1FR2011 has a bulge around the middle due to Phe13 (**Fig. 5B**). The Vpu-O and -P types differ from the M-type in a number of ways (**Fig. 5C-E**). The O-ANT70 and P-RBF168 Vpu have a bulky, positively charged Arg near the N-terminus. Both the size and the charge on this residue give these peptides a different topology, and this could be one of the factors responsible for their inability to bind BST-2. The O-MVP5180 Vpu lacks this Arg residue, but also shows the same activity towards BST-2 as Vpu from O-ANT70 and P-RBF168. The TM substitution, which does not exhibit anti-BST-2 activity, differs remarkably in shape from M-NL4-3 around residue 22 (**Fig. 5F**). While M-NL4-3 has a bulky Trp at this position, the TM substitution has an inward protrusion due to the occurrence of Ala. The SASA profiles of the two peptides also differ around residues 14 and 18. The M-NL4-3 Vpu and F1/F2, on the other hand, show remarkable similarity in their SASA profiles, indicative of similar shapes of the two proteins, except for the bulky Trp22 in F1 (**Fig. 5G-H**). Since the SASA profiles of the BST-2-active peptides are closer to M-NL4-3 than the inactive ones, shape appears to be an important factor in determining the anti-BST-2 activity of these peptides.

**Figure 5.**
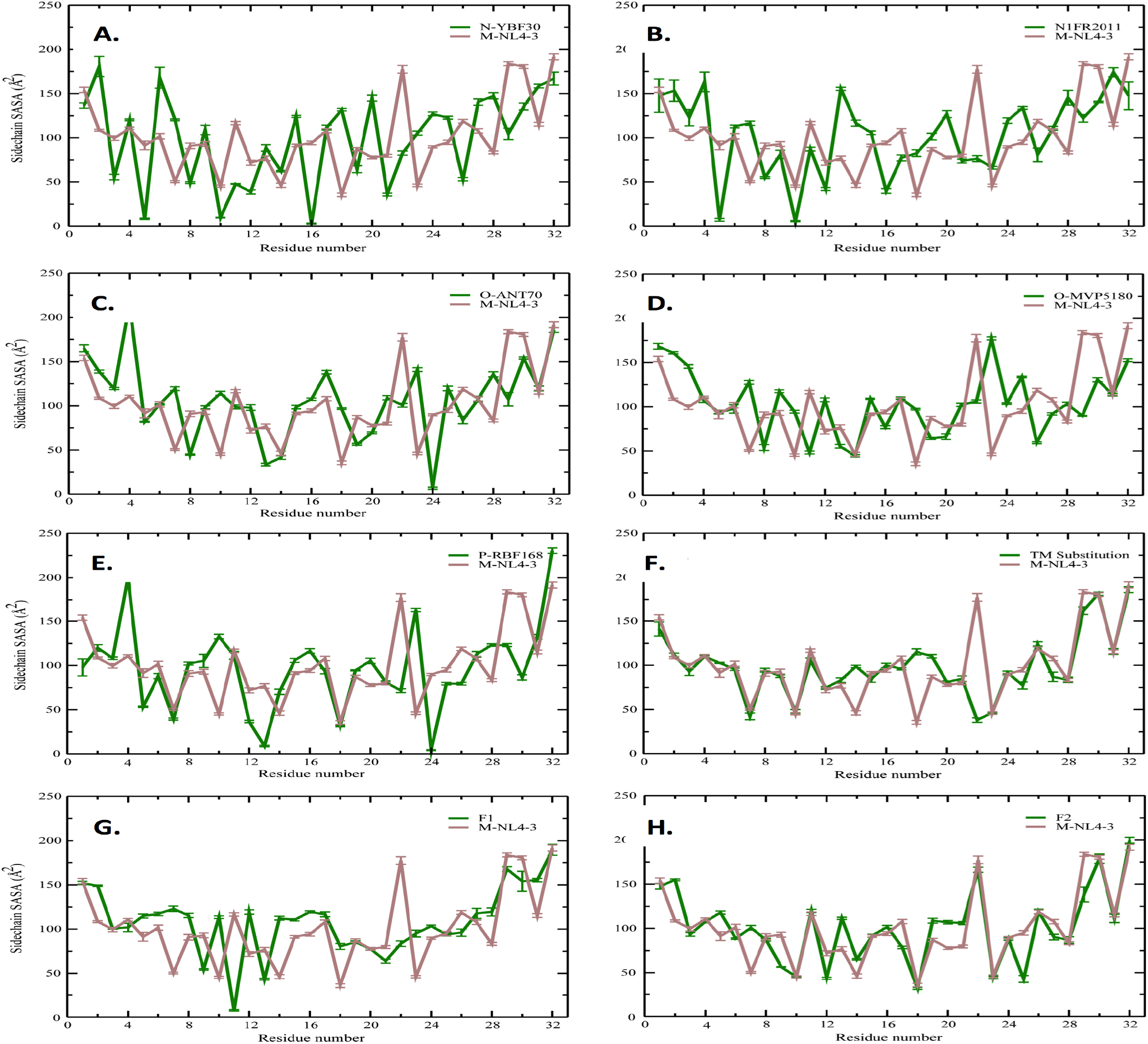
Solvent accessible surface area (SASA) profiles of the Vpu TMDs. The residue-wise values shown are for an average structure over the last 5 ns of simulation in an implicit membrane environment. The SASA of Vpu-M is shown (in brown) for comparison. (A) N-YBF30; (B) N1FR2011; (C) O-ANT70; (D) O-MVP5180; (E) P-RBF168; (F) TM Substitution; (G) F1; and (H) F2.

### Functional effects of recombinant Vpu on cell surface proteins

To test the functional effects of recombinant Vpu proteins on BST-2, we cotransfected HEK293T cells with expression vectors for Vpu-M-EGFP, F1-EGFP or F2-EGFP and BST2-HA, and the cell lysates were Western blotted for the equilibrium levels of these proteins. Compared to an EGFP control, the levels of BST-2 were significantly lower in cells co-expressing Vpu-EGFP, F1-EGFP or F2-EGFP (**Fig. 6A**). While the F2-EGFP levels were also low, it was nevertheless competent at reducing BST-2 levels as efficiently as Vpu-EGFP or F1-EGFP. We then asked whether BST-2 degradation was through the proteosomal pathways by repeating this experiment in the presence of MG132, an inhibitor of proteosomal degradation. There was a large increase in the cellular levels of BST-2, thus confirming its Vpu-mediated degradation to be through the proteosomal pathway (**Fig. 6B**). The Vpu proteins, especially F2-EGFP, were also stabilized following MG132 treatment **(Fig. 6B**). We then analyzed the effects of F1 and F2 on CD4 and BST-2, two cell surface proteins that are important for HIV pathogenesis and are targeted by Vpu. For this, we transfected TZM-bl cells with expression vectors for Vpu-M-EGFP, F1-EGFP or F2-EGFP and quantified cells expressing the Vpu proteins (or EGFP) for surface levels of CD4 or BST-2 by flow cytometry. This showed Vpu-M as well as the F1 and F2 recombinant proteins to significantly reduce the surface levels of CD4 and BST-2 (**Fig. 6C**). The same effect was also observed in Jurkat human CD4+ T cells (**Fig. 6C**). Thus, the F1 and F2 recombinant Vpu proteins are functionally competent at reducing CD4 and BST-2 levels in a cell-type independent manner.

**Figure 6.**
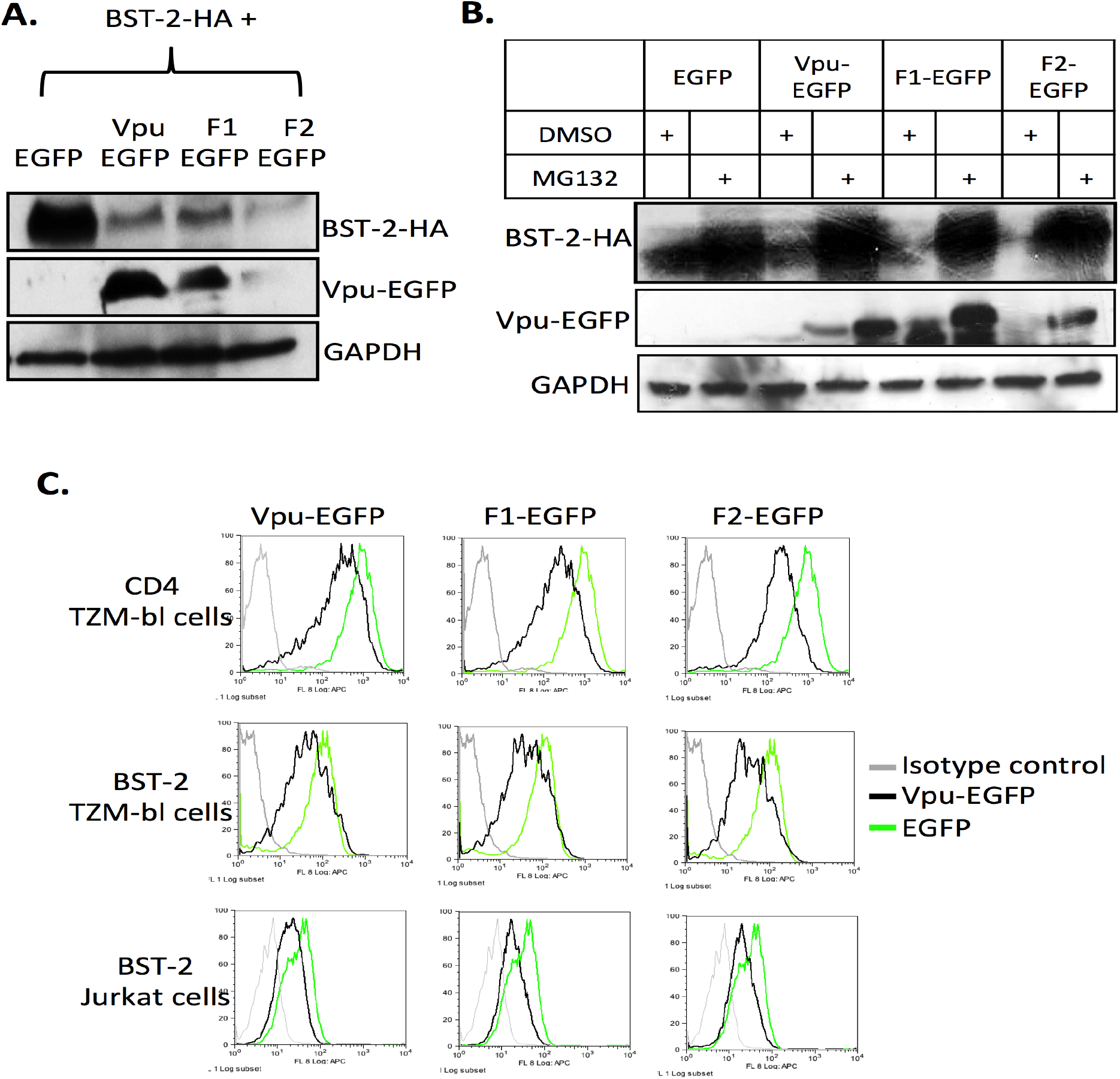
Degradation of BST-2 by Vpu. (A) HEK293T cells were transfected with expression vectors for BST-2-HA (0.5 μg) and EGFP, Vpu-M, F1 or F2 (2.0 μg). After 48 hr, the cell lysates were Western blotted for BST-2 (with anti-HA antibodies) and Vpu (with anti-GFP antibodies); GAPDH served as a loading control. (B) HEK293T cells were transfected as in (A) and either treated with proteosomal inhibitor MG132 (10 μg) or DMSO (vehicle) for 24 hr. The cell lysates were then Western blotted for BST-2 (with anti-HA antibodies) and Vpu (with anti-GFP antibodies); GAPDH served as a loading control. (C) TZM-bl or Jurkat cells were transfected with EGFP-tagged Vpu-M, F1 or F2. After 48 hr cells were surface stained with CD4, BST-2 or isotype control antibodies. The histograms are shown for EGFP-positive cells analyzed using flow cytometry and the FlowJo software.

### Recombinant Vpu inhibits virion production

Since F1 and F2 were functionally competent at reducing BST-2 and CD4 levels, they were also expected to enhance virion production and release. To test this, we transfected HEK293T cells with pNL4-3-EGFP or pNL4-3ΔVpu-EGFP with or without the BST-2 expression plasmid. The pNL4-3ΔVpu-EGFP transfections were also with vectors that expressed Vpu-M, F1 or F2. In cells transfected with pNL4-3-EGFP, there was efficient virus release irrespective of whether BST-2 was co-expressed or not, as evidenced from Western blotting for p24 or TZM-bl infectivity assays for culture supernatants (**Fig. 7A; lanes 1, 2**). In cells transfected with pNL4-3ΔVpu-EGFP, virus was released in the absence but not in the presence of BST-2 (**Fig. 7A; lanes 3, 4**). When these cells were supplemented with Vpu proteins in trans, virus was released only with Vpu-M but not with F1 or F2 (**Fig. 7A; lanes 5-7**). As expected, BST-2 was detectable only in cells that expressed no Vpu-M, F1 or F2 (**Fig. 7A; lane 4**). Compared to cells that expressed Vpu-M, the intracellular levels of p55 and p24 were reduced significantly in cells that expressed F1 or F2. Thus, while the F1 and F2 proteins efficiently degraded BST-2, they also inhibited virion production. In the next set of experiments we evaluated virion production and release and the levels of another viral protein (Nef) at different times post-transfection. Once again, F1 and F2 inhibited the expression of Gag (p55/p24) as well as Nef (**Fig. 7B**). These results confirm the inhibitory effects of F1 and F2, and suggest a possible role for the Vpu TMD in virion production.

**Fig 7.**
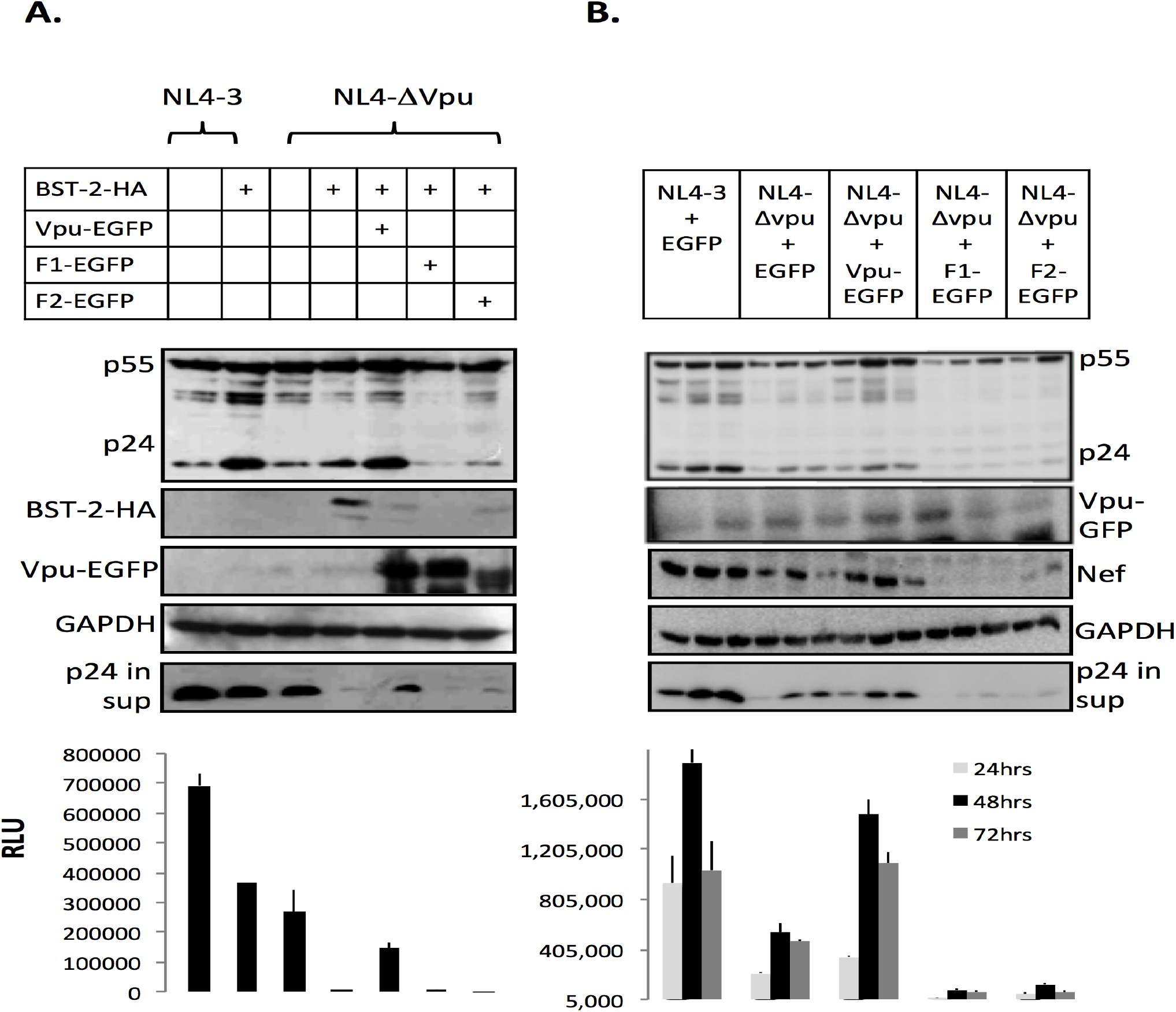
Effect of Vpu on virus production and release. (A) HEK293T cells were co-transfected with the BST-2-HA expression vector (120 ng) and either pNL4-3-IRES-GFP (2 μg) or EGFP control vector (1 μg) to achieve a 1:1 molar ratio; similarly, the cells were co-transfected with BST-2-HA expression vector (120 ng), pNL-ΔVpu-IRES-GFP (2 μg) (or EGFP control) and vectors for EGFP-tagged Vpu-M, F1 or F2 (1 μg) to achieve a 1:1 molar ratio. After 48 hr, cell lysates were Western blotted for Gag (anti-p24), BST-2 by Western blotting the culture supernatant for p24. Virus infectivity was checked in culture supernatants using the TZM-bl assay and expressed as relative light units (RLU). (B) HEK293T cells were cotransfected with pNL4-3-IRES-GFP or pNL-ΔVpu-IRES-GFP (2 μg) and vectors for EGFP-tagged Vpu, F1 or F2 (1 μg) to achieve a 1:1 molar ratio. After 24, 48 or 72 hr, cell lysates and culture supernatants were checked for Gag, Vpu, released virions and virus infectivity as in (A). Additionally, cell lysates were also Western blotted for levels of the HIV-1 Nef protein; GAPDH was used as a loading control.

To test if the F1 and F2 proteins can compete with Vpu-M to limit virion production, we co-transfected HEK293T cells with pNL4-3-EGFP and either F1 or F2 in a 1:1 or 1:2 molar ratio. Virion release was evaluated by Western blotting of the culture supernatants for p24 and TZM-bl assay was used for determining the infectivity of released virions. With increasing F1 and F2, virion release was reduced and this was reflected in the infectivity assay as well (**Fig. 8A**). This suggested that F1 and F2 interfered with Vpu function. We next tested whether F1 and F2 also inhibited virion production across clades by using one HIV-1 subtype B (pNL4-3) and one subtype C (pIndieC1) infectious molecular clone. In both cases, F1 and F2 reduced the intracellular levels of Gag (p55/p24) and Nef, as also the release of new virions (Fig. 8B). We wondered if F1 and F2 rendered Gag susceptible to degradation, and to address this we tested the effects of lysosomal (Leupeptin + Pepstatin A) and proteosomal (MG132) inhibitors on Gag levels. Neither of these inhibitors reversed the effects of F1 and F2 on virus production (**Fig. S2**).

**Figure 8.**
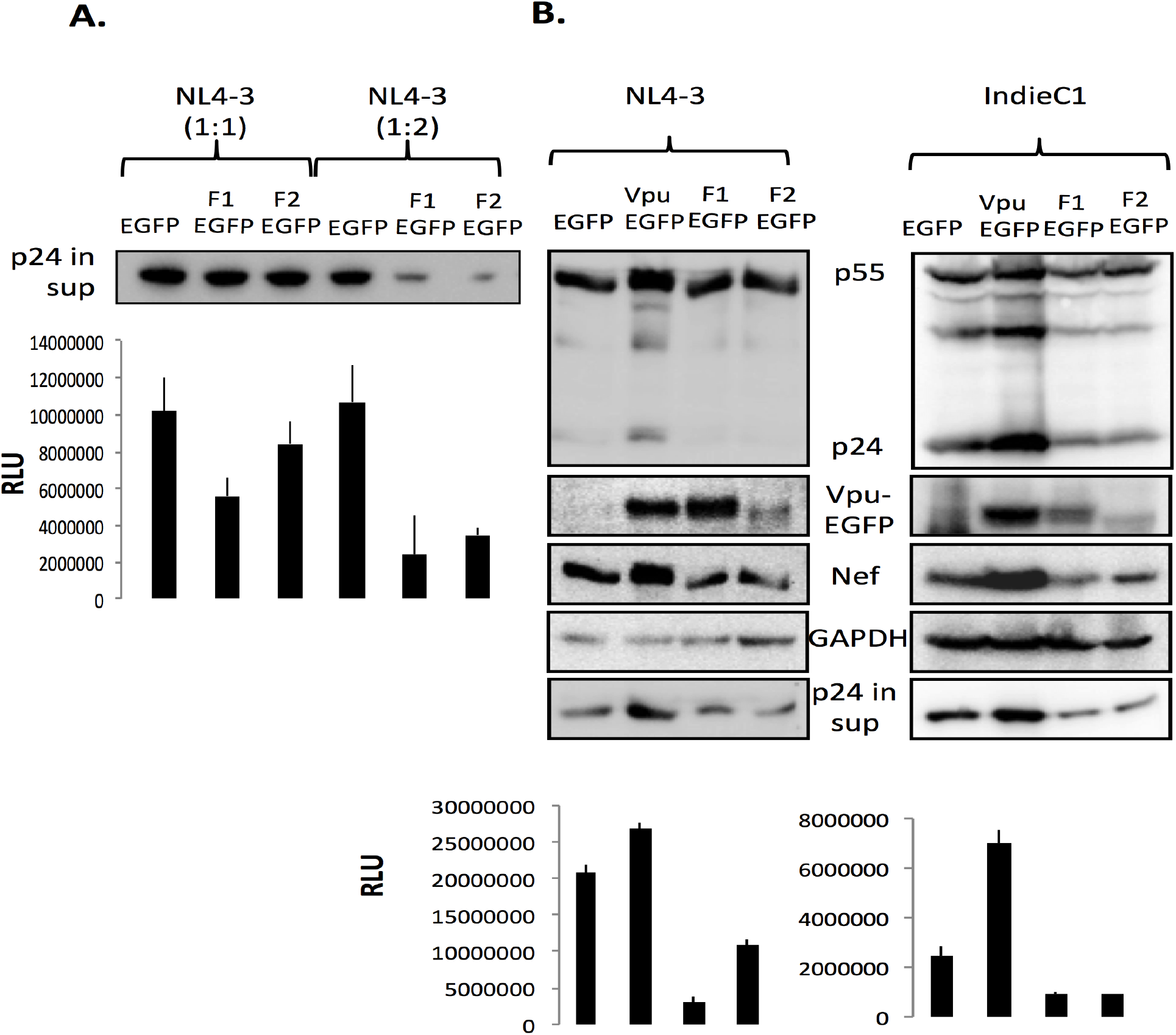
Recombinant Vpu reduces virus production. (A) HEK293T cells were co-transfected with pNL4-3 and either F1-EGFP or F2-EGFP (with EGFP as control) at a 1:1 or 1:2 molar ratio. After 48 hr the virions released in the culture supernatant were estimated by Western blotting with anti-p24 and virus infectivity was quantified with the TZM-bl assay. (B) HEK293T cells were co-transfected with pNL4-3 or pIndieC1 and either Vpu-EGFP, F1-EGFP or F2-EGFP (with EGFP as control) at a 1:2 molar ratio. After 48 hr, cell lysates were Western blotted for Gag, Vpu and Nef proteins; GAPDH served as a loading control. Western blotting the culture supernatant for p24 estimated released virions. Virus infectivity was checked in culture supernatants using the TZM-bl assay.

### The recombinant Vpu proteins inhibit HIV-1 LTR activity

A previous study has shown that Vpu can reduce HIV-1 LTR activity through its effects on NF-kB [46]. We therefore analyzed HIV-1 LTR activity in the presence of Vpu-M, F1 or F2 by co-transfection of HEK293T cells with the Vpu variants (in the background of plasmid pNL4-3 deleted Vpu) and the reporter plasmid pHIV-1-LTR-Luc. There was a significant reduction in LTR activity with F1 and F2 (**Fig. 9A**). The Vpu variants were similarly tested for their effects on LTR activity in the presence of HIV-1 infectious molecular clones pNL4-3 or pADA. Again, there was a significant reduction in LTR activity with F1 and F2 (**Fig. 9B**). To test if this decrease in HIV-1 LTR activity was due to reduced levels of transcription factor(s) known to regulate it, we employed luciferase reporter plasmid transfection assays to check the activities of NF-kB, NFAT and AP1 transcription factors. The results showed no significant differences in the activities of these transcription factors in HEK293T cells expressing Vpu-M-EGFP, F1-EGFP or F2-EGFP (data not shown).

**Figure 9.**
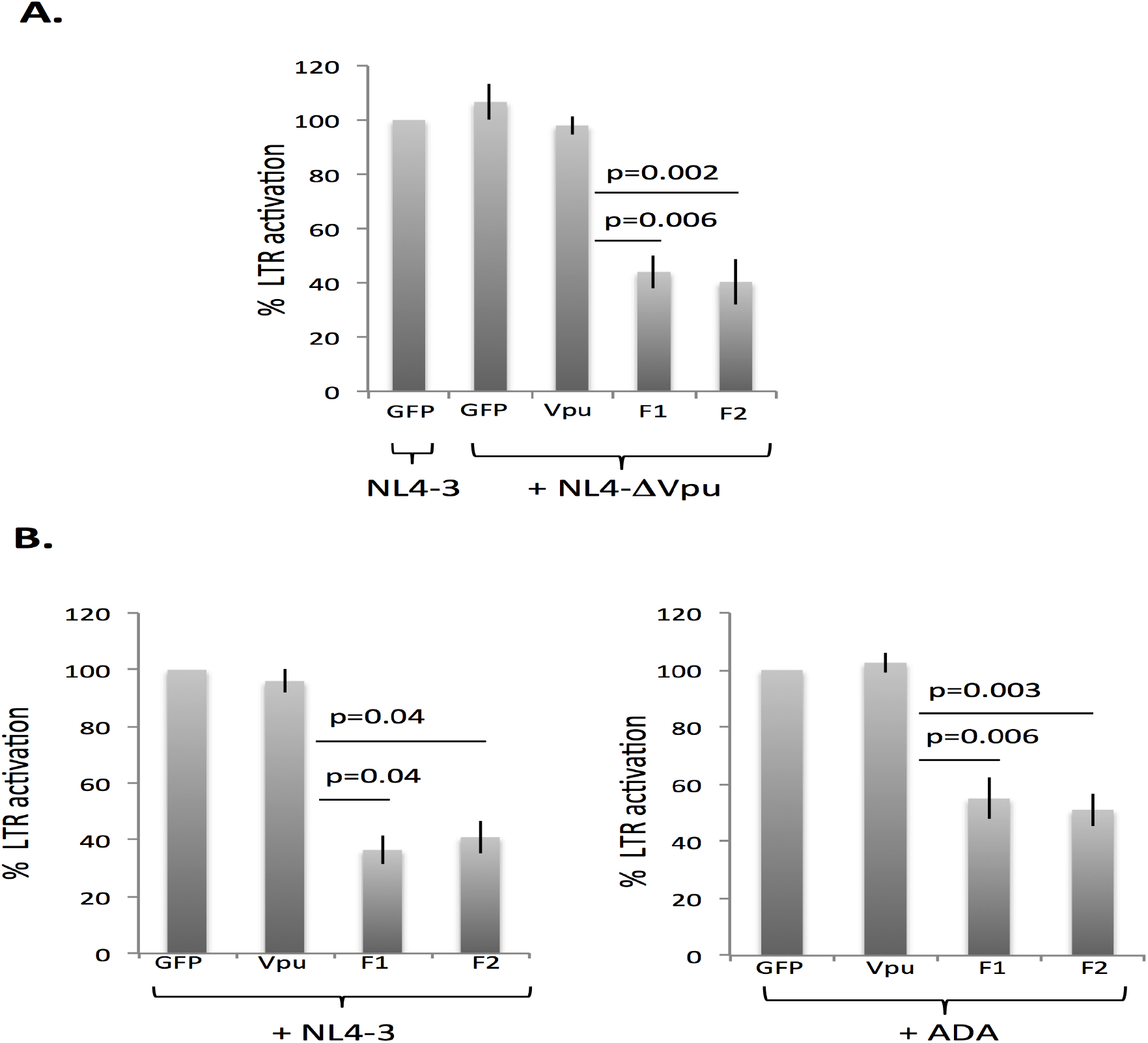
Recombinant Vpu inhibits HIV-1 LTR activity. (A) HEK293T cells were co-transfected with pHIV-LTR-Luc (100 ng) and either pNL4-3-IRES-GFP or pNL-ΔVpu-IRES-GFP (500 ng) and expression vectors for EGFP, Vpu-M, F1 or F2 (250 ng). (B) HEK293T cells were co-transfected with pHIV-LTR-Luc (100 ng) and either pNL4-3 or pADA (500 ng) and expression vectors for EGFP, Vpu, F1 or F2 (500 ng). In both cases, the pRenilla-Luc vector (50 ng) was used to control for transfection efficiency. After 36 hr, cell lysates were prepared and quantified for luciferase expression using the Dual Luciferase System (Promega). The Firefly/Renilla luciferase ratio was taken as a measure of HIV-1 LTR activity, which was normalized to the GFP control taken as 100%. The results are shown as a mean + SD of three independent experiments; the p values are shown.

## Discussion

The Vpu from HIV-1 groups M and N, but not those from groups O and P, overcome the BST-2 host cellular restriction factor and promote the release of newly assembled virions from the surface of infected cells. It has been suggested that BST-2-active Vpu contains the motif A10XXXA14XXXA18 in its transmembrane domain (TMD), and that Vpu types lacking this specific sequence do not possess anti-BST-2 activity [20]. Additionally, Leu11 and N-terminal residues of the Vpu TMD have also been shown to be important for targeting BST-2 [45, 47]. Finally, the anti-BST-2 activity of Vpu is also compromised when the polarity of its TMD is changed even though the key sequences are conserved. The determinants of interaction between the Vpu and BST-2 TMDs and the effects on BST-2 degradation were investigated in this study.

We selected two topological homologs for the TMD of BST-2-active Vpu-M from TOPDB and constructed Vpu variants. These included F1 with TMD of the human Asialoglycoprotein receptor and F2 with TMD of the E. coli Acriflavin protein. Like Vpu-M, both of these variants interacted with BST-2 and reduced its levels through the proteosomal degradation pathway. The Vpu-M/BST-2 interaction is driven by hydrophobic interactions, with nonpolar residues lining the interface between the two proteins. The nonpolar nature of the interactions is also seen in the F1 and F2 recombinants. Although previous studies have shown Ala14 and Trp22 on Vpu-M to be important for its interaction with BST-2 [20], F1 and F2 can interact with BST-2 despite the absence of Trp22 in F2 and Ala14 in both F1 and F2. SASA profiles show that while Vpu-M, F1 and F2 TMDs have similar shape, those of BST-2 inactive Vpu-O and Vpu-P differ in shape. The N-termini of BST-2 active Vpu-M, Vpu-N, F1 and F2 also carry nonpolar residues, unlike Vpu-O and Vpu-P, suggesting that a nonpolar N-terminus is also required for the anti-BST-2 activity of Vpu [45, 47].

The Vpu proximal hinge region (KILRQ) interacts with the small glutamine-rich tetratricopeptide repeat (SGTA) and helps transport Gag to the site of virus assembly [28, 29]. In F1 and F2, the entire TMD was replaced without affecting the proximal hinge region of Vpu. Expectedly, like Vpu-M, F1 and F2 also interacted with SGTA in co-immunoprecipitation assays (data not shown), indicating that the function of Vpu proximal hinge region is not affected by TMD replacement. Despite full SGTA interaction activity, and thus their ability to transport Gag to assembly sites, virus release was compromised in the presence of F1 and F2.

Even in the absence of BST-2, the F1 and F2 do not support virus release, and intracellular Gag levels were reduced in these cells compared to Vpu-M. This was not rescued with inhibitors of the proteosomal and lysosomal degradation pathways, suggesting that the effect of the F1 and F2 variants was upstream of intracellular protein expression. The F1 and F2 variants reduced HIV-1 LTR activity by 50-60%, but this could not be linked to a corresponding decrease in three major transcription factors that affect HIV-1 LTR activity. The cellular Twik-related Acid Sensitive K+ channel 1 (TASK-1) and HIV-1 Vpu proteins suppress transcription of unintegrated HIV-1 DNA through an NF-kB-dependent mechanism [46]. Thus, with Vpu trans-complementation, viral protein levels should decrease. However, this was not the case in our study as well as in other studies [3, 20, 35, 48]. Despite the presence of homologous TMDs, Vpu variants reduce viral protein levels by decreasing LTR activity even when the activity of transcription factors like NF-kB and NFAT was not altered. Emeagwali et al. also showed that TASK-1, which is a structural homolog of the Vpu-TMD with some identical amino acids [49], suppresses HIV-1 transcription. Although F1 and F2 possess anti-BST-2 activity, they do not support virus replication. Furthermore, F1 and F2 also suppress the activity of Vpu resulting in the inhibition of virus production in subtype B (NL4-3) as well as subtype C (IndieC1) backgrounds.

In conclusion, topology is a critical determinant of the interaction between BST-2 and Vpu, with shape and the occurrence of a hydrophobic interface being important factors. Vpu from the M group of HIV-1 is therefore topologically adapted for enhancing virus release via BST-2 downregulation. However, its topological homologs F1 and F2 suppressed HIV-1 LTR through a mechanism that is independent of the modulation of three major transcription factors – NF-kB, NFAT and AP1. The pathway might involve histone deacetylase (HDAC) or zinc finger proteins, which also have effects on HIV-1 LTR activity [50–52]. These will be tested in future studies.

## Materials and Methods

### Identification of Vpu homologs

To identify helical transmembrane proteins with TMDs homologous to that of Vpu, the sequences of all alpha bitopic and alpha polytopic proteins were extracted from the Topology Data Bank of Transmembrane Proteins (TOPDB) [36]. Transmembrane segments were also extracted from the Membrane Protein Topology Database (MPtopo) [37]. Pairwise alignment was performed between the TMD of Vpu-M (VAIVALVVAIIIAIVVWSI) and all sequences obtained from the two databases using EMBOSS v6.6.0 [53] with the Needleman-Wunsch algorithm and the BLOSUM45 matrix. Sequences with similarity greater than 70% and a score higher than 25.0 were selected, and two of these were chosen as potential topological homologs for further investigation.

### Plasmids and Expression Vectors

The Vpu-EGFP construct was prepared using the vpu gene from the NL4-3 genome. The TMDs of Vpu were replaced with those from the human asialoglycoprotein and E. coli Acriflavin proteins to give F1-EGFP and F2-EGFP, respectively. Oligonucleotides were synthesized for the amino acid sequences MQPILLLLSLGLSLLLLVVVCVIIVII (F1) and MQPIITIVSAMALSVLVALILWSIVII (F2). These were the following: F1-Forward, AATTCATGCAACCTATACTGCTGCTGC TGAGCCTGGGCCTGAGCCTGCTGCTGCTGGTGGTGGTGTGCGTGATC; F1-Reverse, GATCACGCACACCACCACCAAGCAGCAGCAGGCTCAGGCCCAGGCTCAGCAGCA GCAGTATAGGTTGCATG; F2-Forward, AATTCATGCAACCTATAATCACCATCGTGAGCGCCATGGCCCTGAGCGTGCTGGTGGCCCTGATCCTG; and F2-Reverse, CAGGA TCAGGGCCACCAGCACGCTCAGGGCCATGGCGCTCACGATGGTGATTATAGGTT GCATG. The forward and reverse oligonucleotides were annealed and inserted in the Vpu-EGFP vector, which was PCR amplified with oligonucleotides: VpuF1, GTGTGGTCCAT AGTAATCATAGAATATAGGAAAATA or VpuF2, TGGTCCATAGTAATCATAGAAT ATAGGAAAATATTAAG; and VpuR, CATGAATTCGAAGCTTGAGCTCGA. The PCR products were digested with restriction enzymes EcoRI and BamH1 and the annealed F1 and F2 fragments ligated into these vectors. The plasmids pNL4-3-IRES-GFP and pNL-ΔVpu-IRES-GFP and the expression vector for BST-2-HA were obtained from Dr. Frank Kirchhoff (Ulm University, Germany), and the expression vectors for GFP-tagged Vpu-N, -O and -P were from Dr. Patricia Cannon (University of Southern California, USA).

### Cell culture and transfection

HEK293T and TZM-bl cells (HIV indicator cell line; NIH AIDS Reagent Program) were cultured in DMEM with 10% fetal calf serum (FCS), 10 unit/ml Penicillin and 10 ug/ml Streptomycin (Invitrogen). Transfections were performed using jetPrime reagent (Polyplus, Germany) following the manufacturer’s protocol.

### Confocal Imaging

For cell membrane association, TZM-bl cells were transfected with empty vector or those expressing Vpu-M, F1 or F2. After 48 hr, the cells were stained with 1:1000 dilution of the PKH26GL dye (Sigma Aldrich) for 2 min at room temperature. For Vpu and BST-2 co-localization, TZM-bl cells were transfected with empty vector or those expressing Vpu-M, F1 or F2. After 48 hr, the cells were fixed with 4% paraformaldehyde and permeabilized with 0.4% Triton X-100 for 10 min each. These were then stained with 1:100 dilution of mouse anti-BST-2 antibody (Abcam) for 1 hr followed by 1:500 dilution of anti-mouse Alexa-594 (Calbiochem) for 45 min at room temperature. All images were acquired using a Nikon A1R confocal microscope at 60x magnification.

### BST-2/Vpu interaction

HEK293T cells were co-transfected with expression vectors for BST-2-HA and EGFP-tagged Vpu-M, F1 or F2. The cells were lysed in lysis buffer on ice, centrifuged at 16,000xg for 10 min and the clarified lysates were pre-adsorbed to Protein A or Protein G beads (GE Healthcare Bioscience) for 2 hr at room temperature. The lysates were then immunoprecipitated overnight with rabbit anti-HA antibody (for BST-2) or mouse anti-GFP antibody (for Vpu-M, F1 and F2) at 4° C with rotation, followed by Protein A or Protein G beads for 2 hr at room temperature. The beads were then boiled in SDS-dye loading buffer and the supernatants subjected to Western blotting with mouse anti-GFP antibody or anti-rabbit HA antibody to detect co-precipitated Vpu-M/F1/F2 and BST-2, respectively.

### Flow cytometry

To analyze BST-2 cell surface levels, TZM-bl cells were transfected with pEGFP-N1 vector or expression vectors for EGFP-tagged Vpu-M, F1 or F2 using the jetPrime transfection reagent. Jurkat cells were transfected with these plasmids using a nucleofection kit (Lonza). The cells were harvested 48 hr post-transfection with 0.02% EDTA and resuspended in 1X PBS containing 0.01% FCS (stain buffer). The transfected cells were stained with 1:100 dilution of anti-mouse BST-2 (Abcam) or isotype control antibody (R&D system) on ice, washed three times with stain buffer, stained with 1:600 dilution of secondary anti-mouse Alexa 647 (Calbiochem), and then washed three times. The cells were then acquired on a DAKO Cyan ADP flow cytometer, and BST-2 surface staining analyzed on cells gated for GFP expression using the Flow Jo software.

For CD4 surface levels, TZM-bl cells were similarly transfected followed by staining with 1:50 dilution of anti-mouse monoclonal CD4-APC conjugated antibody (Abcam) or isotype control (Abcam) on ice for 45 min. The cells were washed three times with stain buffer, acquired and analyzed as above.

### Virus replication and release

HEK293T cells were cotransfected with either pNL4-3 or pNL-ΔVpu-IRES-GFP, together with expression vectors for EGFP (control) or EGFP-tagged Vpu, F1 or F2, and BST-2-HA After 48 hr cells were harvested in lysis buffer containing 150 mM NaCl, 20 mM Tris pH 7.5, 1 mM EDTA, 1% Triton X-100, 1 mM EGTA, 2.5 mM Sodium pyrophosphate, 1 mM Na3VO4 and protease inhibitor cocktail (Roche, Germany). After centrifugation at 16,000xg for 10 min, the protein concentrations in clarified lysates were estimated using the Bradford assay (Bio-Rad). The culture supernatants were collected and filtered through a 0.4 μm membrane filter. The cell lysates were subjected to Western blotting for viral proteins (Gag, Nef); the culture supernatants were subjected to Western blotting for p24 and ELISA for quantification of released viral particles. Virus infectivity was quantified using the HIV indicator TZM-bl cell line as previously described [54]. For this, 10,000 TZM-bl cells per well were seeded in 96-well plates (Nunc-Greiner). The following day cells were infected with an equal volume of culture supernatant for 4 hr. After 48 hr, cells were lysed on the plate, used for the Luciferase reporter assay (Steady Glow Luciferase Kit, Promega) and quantified as Relative Light Unit (RLU) on a Luminometer (GLOMax 20/20). Similar experiments were also carried out with infectious molecular clones for another subtype B HIV-1 (pADA) and a subtype C HIV-1 (pIndieC1).

### Western blot

Samples containing equal amounts of protein were subjected to electrophoresis on SDS-12% polyacrylamide gels, and then transferred to 0.4 μm nitrocellulose membranes (Millipore). The membrane was blocked with PBS-0.01% Tween-20 (PBST) containing 5% nonfat dry milk for 1 hr at room temperature, and then washed thrice with PBST. The washed membrane was incubated with appropriately diluted primary antibodies, washed and incubated with secondary antibody conjugates. The following primary antibodies were used – p24 hybridoma (NIH AIDS Reagent Program), GFP antibody (Santa Cruz) for EGFP-tagged Vpu/F1/F2 detection, Nef antibody (EEH1, NIH AIDS Reagent Program) and anti-HA antibody for HA-tagged BST-2 protein (Santa Cruz). Secondary antibodies conjugated to HRP (Calbiochem) were used at appropriate dilution, the blots were developed using the ECL Reagent (Santa Cruz) and images were captured on a FluorChem M machine (Protein Simple, Santa Clara, CA, USA).

### Equilibration of Vpu, F1, F2 and BST-2

The sequences of the TMDs of the peptides used in the study are shown in Table 2. As described previously [55, 56], extra residues have been considered on both sides of the TMD to ensure that there are no destabilizing effects due to abrupt termination of the TMD. The peptides were modeled as idealized a-helices using SYBYL7.2 [57]. The TMDs were then simulated in an implicit membrane environment using CHARMM [58] with the CHARMM22 protein force field with CMAP corrections [59, 60]. The NMR structure with PDB ID 2LK9 was used for modeling the BST-2 TM domain [34]. Simulations were carried out in a solvated membrane environment using the NAMD program [61] with the CHARMM22 protein force field with CMAP corrections [59, 60], the CHARMM36 lipid force field [62], the TIP3P water model [63], and optimized parameters for K^+^ and Cl^−^ [64]. All simulation details are described in Supporting Information.

**Table 2.**
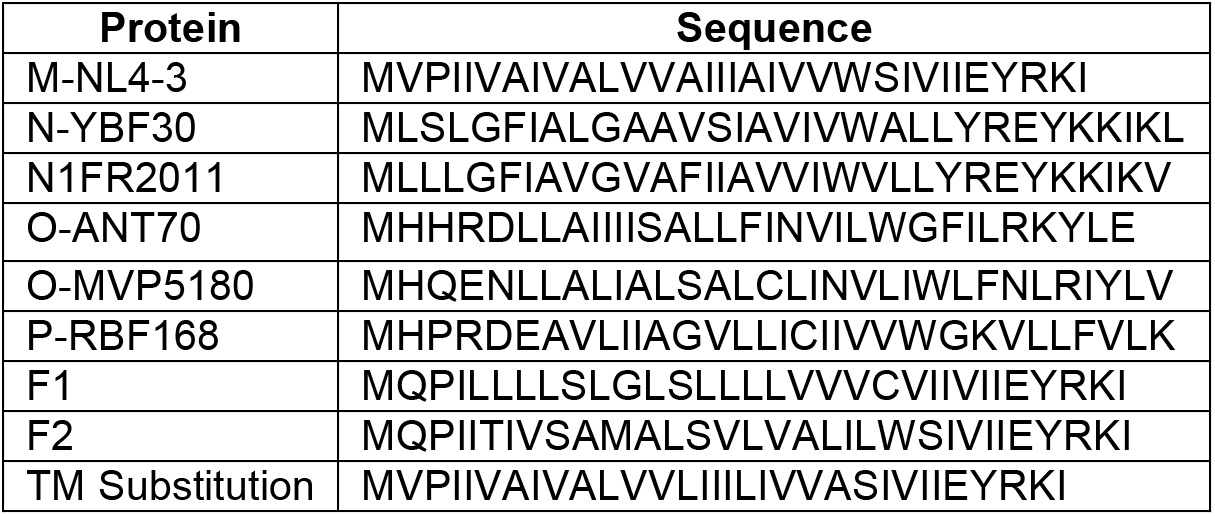
Sequences of the TMDs of Vpu from different HIV-1 groups and the topological homologs used in this study

### BST-2/Vpu docking studies

The equilibrated BST-2 structure was docked on to each of the equilibrated wild type and recombinant Vpu peptides using the HADDOCK server [65]. All residues excluding the terminal 6 residues in BST-2 and the peptides were regarded as active residues in the docking, and amino acids surrounding the active residues were regarded as passive residues. For further analysis, representative docked structures were selected from the cluster (the HADDOCK program generates clusters of docked complexes) with the most favored HADDOCK score. A representative structure was accepted only if it satisfied the following criteria: (1) the two docked peptides were in an anti-parallel orientation, and (2) the two helices were aligned such that both would fit into a membrane [see Results section for details]. If a complex did not satisfy either of the above criteria, a complex from the second most favorable cluster was selected. The complex was set up in a fully solvated membrane environment, and equilibration was carried out using the same parameters as those used for unbound BST-2. Production runs were carried out in the NPT ensemble for 10 ns with a time-step of 2 fs.

### Sequence and topology analysis of the peptides

The Clustal Omega server [66] was used to perform a multiple sequence alignment and cluster the sequences. Curvograms of the clusters were plotted using the TreePlot Server [www.bioinformatics.nl/tools/plottree]. The topology analysis on the wild type and recombinant Vpu peptides was performed on an average structure over the last 5 ns of the equilibration runs on the peptides.

### Measurement of HIV-1 LTR activity

HEK293T cells were co-transfected with the following plasmids: (1) pHIV-LTR-Luc (100 ng), (2) pRenilla-Luc (50 ng), and (3) either pNL4-3-IRES-GFP, pADA or pNL-ΔVpu-IRES-GFP (500 ng) plus EGFP, Vpu-EGFP, F1-EGFP or F2-EGFP (250 ng). The pRenilla-Luc vector was used as a control for transfection efficiency. After 36 hr, Firefly and Renilla luciferase activities were quantified. The Firefly/Renilla luciferase ratio was taken as a measure of HIV-LTR activity.

## Acknowledgements

We are grateful to Dr. Frank Kirchhoff (Ulm University, Germany) for providing pNL4-3-IRES-GFP/pNL-ΔVpu-IRES-GFP and BST-2-HA plasmids, and to Dr. Patricia M. Cannon (University of Southern California, USA) for providing GFP tagged-Vpu M, N, O and P subtype plasmids. This work was supported by a grant to SJ from the Department of Biotechnology and to UDP from the Department of Biotechnology. NK received a Senior Research Fellowship from the Indian Council of Medical Research; PP from the Council of Scientific and Industrial Research, India; and SP from the Department of Biotechnology.

## Supporting information

**Figure S1.**
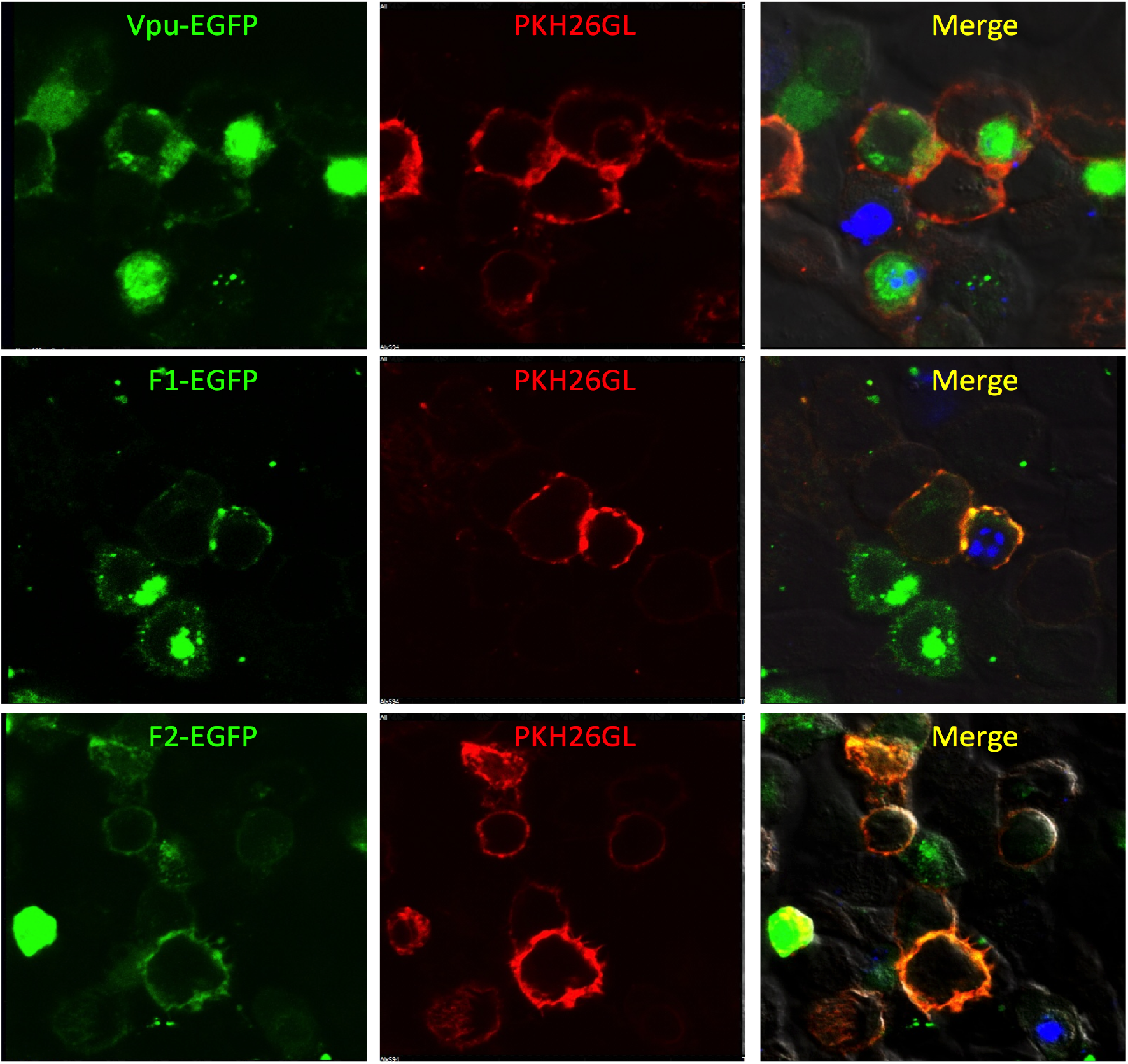
Localization of wild-type and rec-Vpu in the cell membrane. TZM-bl cells were transfected with Vpu or rec-Vpu-GFP by Jetprime reagent. 48 hrs post-transfection, cells were stained with cell membrane tracker PKH26GL for 2-3 minutes. Cell images were captured with confocal imaging. Vpu or Rec-Vpu are seen to localize in the cell membrane, as can be seen from the merged images.

**Figure S2.**
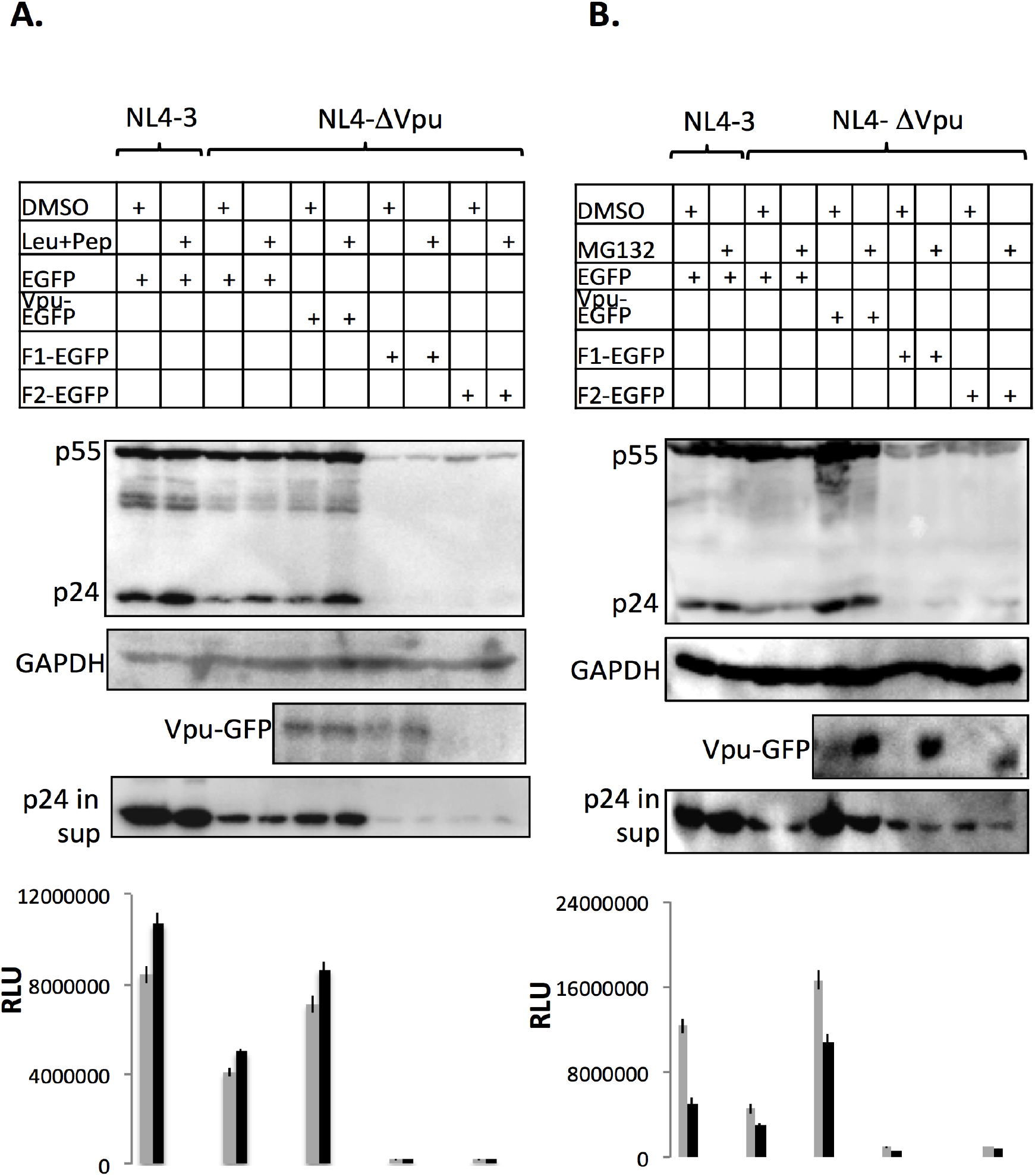
Gag protein degradation and proteosomal/lysosomal pathways. 293T cells were cotransfected with NL-43-IRES-GFP, NL-ΔVpu-IRES-GFP/ EGFP or Vpu/rec-Vpu-GFP (1:1) molar ratio. 24 hrs post-transfection, cells were treated with MG132 proteosomal inhibitor (10μM) or lysosomal inhibitors leupeptin and pepstatin (40μM) for 24 hrs. The cells were then harvested for making cell lysates using lysis buffer. p24 levels were then detected inside the cells or in supernatant by western blot analysis. p24 levels could not be enhanced even with proteosomal/lysosomal inhibitor treatment, indicating that p24 protein decline is not due to proteosomal/lysosomal pathways.

## Supporting Information

### Simulation details for equilibration of wild-type and recombinant Vpu

The TMDs of the different Vpu types were modeled using the program SYBYL7.2 (Tripos International, St. Louis, Missouri, USA; http://www.tripos.com). The helices were then placed in an implicit membrane environment with hydrophobic thickness 33 Ǻ and aligned along the normal to the membrane. The Generalized Born model with a simple SWitching function (GBSW) module was used with a surface tension coefficient of 0.03 kcal mol^−1^ Ǻ^−2^ [1, 2]. Nonbonded interactions were treated by using a smoothing function between 8 Ǻ and 10 Ǻ, and updating nonbonded lists with a cutoff of 12 Ǻ. Simulations were carried out for a total of 10 ns using the CHARMM program [3, 4] with the CHARMM22 all-atom protein force field including CMAP corrections [5, 6]. An integration time-step of 1 fs was used, and the temperature was maintained at 298 K using the Nose-Hoover thermostat [7, 8].

### Simulation details for equilibration of BST-2

The starting conformation for modeling the BST-2 TMD was the NMR structure with PDB ID 2LK9 [9]. Missing side chains were added using the SCWRL program [10]. To avoid end effects, additional residues were added to the two termini using the molecular visualization program UCSF Chimera [11]. On the whole, the model included residues 18 to 49 (KRCKLLLGIGILVLLIIVILGVPLIIFTIKAN). The protein was then placed in a fully solvated membrane environment using the CHARMM-GUI Membrane Builder [12, 13]. A lipid bilayer of 1-palmitoyl-2-oleoyl-sn-glycero-3-phosphocholine (POPC) was set up around the protein, and bulk water of 15Ǻ thickness was placed above and below the bilayer. Potassium and chloride ions were added to attain a concentration of 0.15 M, which is close to physiological salt concentrations.

The simulations were performed with the NAMD program [14, 15] using the CHARMM22 protein force field with CMAP corrections [5, 6], the CHARMM36 lipid force field [16], the TIP3P water model [17], and optimized parameters for K^+^ and Cl^−^ [18]. Periodic boundary conditions were employed, and long-range electrostatics was treated using the particle mesh Ewald method [19]. Nonbonded interactions were cut off at 12 Ǻ with a switching distance of 10 Ǻ. Langevin dynamics was used to maintain a constant temperature of 303.15 K [14]. The six-step equilibration scheme proposed by Jo and coworkers was used [12]. Briefly, positional harmonic restraints were applied on the protein backbone, protein sidechain, and ions. Restraints were also used to prevent water molecules from entering the hydrophobic core of the membrane, and to keep the head groups of all lipid molecules in the same plane. The values of the restraints were gradually reduced to zero over the six steps of equilibration. The first two steps of equilibration were carried out in the NVT ensemble, and the last four steps in the NPT ensemble. The total equilibration time was 675 ps. Production runs were carried out in the NPT ensemble for 5 ns with an integration time-step of 2 fs.

